# How many dinosaur species were there? Fossil bias and true richness estimated using a Poisson sampling model (TRiPS)

**DOI:** 10.1101/025940

**Authors:** Jostein Starrfelt, Lee Hsiang Liow

## Abstract

The fossil record is a rich source of information about biological diversity in the past. However, the fossil record is not only incomplete but has inherent biases due to geological, physical, chemical and biological factors. Our knowledge of past life is also biased because of differences in academic and amateur interests and sampling efforts. As a result, not all individuals or species that lived in the past are equally likely to be discovered at any point in time or space. To reconstruct temporal dynamics of diversity using the fossil record, biased sampling must be explicitly taken into account. Here, we introduce an approach that utilizes the variation in the number of times each species is observed in the fossil record to estimate both sampling bias and true richness. We term our technique TRiPS (<u>T</u>rue <u>R</u>ichness estimated using a <u>P</u>oisson <u>S</u>ampling model) and explore its robustness to violation of its assumptions via simulations. We then venture to estimate sampling bias and absolute species richness of dinosaurs in the geological stages of the Mesozoic. Using TRiPS, we estimate that 1936 (1543-2468) species of dinosaurs roamed the Earth during the Mesozoic. We also present improved estimates of species richness trajectories of the three major dinosaur clades; the sauropodomorphs, ornithischians and theropods, casting doubt on the Jurassic-Cretaceous extinction event and demonstrating that all dinosaur groups are subject to considerable sampling bias throughout the Mesozoic.

## Introduction

One of the main goals of palaeobiology is to reconstruct diversity using information from the fossil record. While the patterns of diversity in space and through time are interesting in themselves, understanding the dynamics of taxon richness is also the first step in elucidating the biotic and abiotic forces that shape the spatial and temporal variation in taxon diversity. In other words, we need an accurate picture of patterns of past diversity to understand processes that operate on long time scales. As in all study systems where data samples in themselves cannot be assumed to represent a complete picture of the underlying population, richness studies based on the fossil record must consider the incompleteness of the fossil record.

Not all organisms enter the fossil record or have the same potential of doing so. Once created, a fossil record (a physical record of the existence of organisms that were alive in the past) is subject to eternal loss through physical processes such as erosion. Whether or not a fossilized organism can be found is also affected by variability in outcrop accessibility. Last but not least, sampling intensity encompassing factors including academic/commercial interest, geographic location and sampling design also influence information from the fossil record we have access to. While some of these factors contribute to noise in our inference of historical patterns and processes, and thus only cloud biological signals, others may cause systematic bias so as to yield misleading results if the data are interpreted at face value or with inappropriate methods.

Several classes of approaches for estimating richness using an incomplete fossil record have been developed. These might be loosely grouped into subsampling approaches, phylogenetic corrections and residual approaches. It is not our purpose to give a full overview of the approaches available, which have variously been reviewed elsewhere (see e.g. 1,2), but we briefly describe these in order to clarify why we have developed a new approach here. Subsampling approaches, including rarefaction (reviewed in 1) and Shareholder Quorum Subsampling (SQS; 3,4), attempt to standardize temporal (or spatial) samples so as to achieve comparable relative richness across samples. Phylogenetic approaches use phylogenetic hypotheses of the clade in question to infer ghost lineages unobserved in the fossil record but that must have existed as implied by the given phylogenetic hypothesis (5). These ghost lineages are thus assumed to give a minimum estimate of the number of lineages we have failed to observe in the fossil record. The residual approach (6–9) assumes that a given proxy for sampling (e.g. outcrop area or number of fossil bearing collections) captures the biases that might influence our observations and uses the proxy to model how a signal driven entirely by sampling would appear. Deviations from such a model are thought to reveal the real troughs and peaks in richness. In all of these approaches, we can only hope to estimate relative richness through time and not true richness. Additionally, none of these approaches attempts to estimate the bias itself, i.e. the differential sampling across time, space or taxa. Without an estimate of sampling bias that is separate from richness estimates, it is difficult to shed light on the Common Cause Hypothesis, which states that a common factor affects both biological dynamics and sampling (10–13).

Here, we introduce an approach that explicitly models the sampling process while estimating richness, using multiple observations of fossils belonging to an organismal group. We named it TRiPS (<u>T</u>rue <u>R</u>ichness estimated using a <u>P</u>oisson <u>S</u>ampling model). While we and others have used the simultaneous estimation of extinction, speciation and sampling processes to study diversification processes (14–17), there has not been a direct attempt to use multiple observations of fossil species to estimate true richness, rather than relative richness, while simultaneously and explicitly estimating sampling, as far as we are aware. Specifically, TRiPS assumes that species observed multiple times in a given time interval have a relatively high probability of fossilization and modern day discovery. We use this type of information across species that are likely to have similar fossilization potential and modern day discovery rates to estimate the number of species we might be missing and hence the true number of species that might have existed.

Dinosaurs are used as an example to illustrate our approach, not least because there is a lot of interest in estimating both the absolute (18–20) and relative temporal richness (21–25) of dinosaur taxa. As earlier analyses suggest that the three major dinosaur groups Sauropodomorpha, Ornithischia and Theropoda exhibit both different diversity dynamics and differential impact of sampling bias (9,25,26), we estimate sampling rates and true richness for all dinosaurs as well as these groups independently. We present stage-specific dinosaur sampling rates (i.e. bias) and dinosaur species richness through the Mesozoic as estimated from TRiPS, compare our estimates with those discussed in the literature and present simulations that explore the power of our approach and the sensitivity of TRiPS to violations of key assumptions.

## Methods and data

### Data

We downloaded records of Dinosauria, Ornithischia, Sauropodomorpha and Theropoda separately from the Paleobiology Database (PaleoDB, https://paleobiodb.org/#/, download August 13^th^ 2015) using the R package paleobioDB (27). Each row of data downloaded from the PaleoDB is associated with an observation of a taxon, its location and age range of the outcrop it was found in, and their metadata. In each download, we first discarded all records that were not identified to the species level by requiring that the ‘matched rank’ of the entry was “species.” Second, we removed all *nomina dubia* and *nomina nuda* by comparing observations with a list of invalid taxa from PaleoDB. Third, we removed ichno- and ootaxa by matching our downloaded data with recently compiled lists of such taxa (28). We further identified four ichno-genera *(Harpedactylus, Yunnanpus, Saurichnium* and *Tetrapodium*), when comparing the download of all dinosaur observations with the subclades. Lastly we required that each observation was temporally assigned to one or more geological stages, i.e. observations only dated to epochs or periods were discarded. The final four datasets had 3122 observations of Dinosauria (974 genera and 1124 species), 1150 observations of Ornithischia (306 genera, 360 species), 602 observations of Sauropodomorpha (215 genera with 246 species) and 1366 observations of Theropoda (451 genera with 516 species).

For each species, we tallied the number of observations per species in each stage in the Mesozoic, generating an observation count matrix. As already noted, the reported age range of a given record can span more than one geological stage. In such cases, we assigned a stage within the reported age range with a probability that is proportional to the duration of those stages. Because of this probabilistic assignment of records to stages within given age ranges, we performed TRiPS analyses (described below) on 100 replicated observation count matrices and used the median estimated sampling rate for species richness estimation. We also analysed genus level data but because both richness and sampling dynamics are similar to species level dynamics, we refer readers to the Electronic Supplementary Material (ESM) for genus level estimates.

### Modelling fossil sampling as a Poisson process

Here, we treat the process of fossil sampling, which we will estimate from records from the PaleoDB (see previous section) as the combined processes of fossilization and detection. In other words, a sampling event is one fossil observation of a particular species, subsuming everything that has to occur from the point an individual died in the past, through fossilization and detection and finally ending up as an observation in a scientific database. Bias in the fossil record is then interpreted as systematic differences in the rate of sampling across time, space or organismal groups.

We assume that sampling can be viewed as a homogenous Poisson process inside a particular time interval. While others have also treated fossil sampling as a Poisson process (e.g. 29–33), ours, to the best of our knowledge, is the first attempt to use such an approach to estimate true richness. Formally, let the Poisson intensity (or rate) *λ_t_* be the parameter controlling the sampling process in a given time interval *t*. The number of observations *O_i,t_* for a species *i* in that time interval *t* with duration *d_t_* has a Poisson distribution with mean *λ_t_d_t_.* The likelihood of the sampling rate *λ_t_* given *O_i,t_* observations in that interval is then

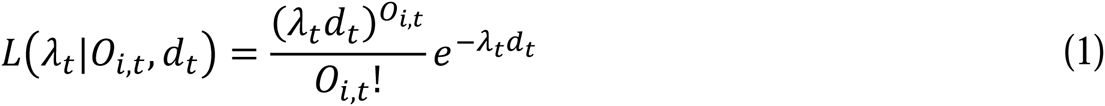

Here we explicitly assume that a species detected in a time interval is extant during that whole time interval. Because any species that is represented in the database must have left at least one detected fossil we must condition the likelihood of *λ_t_*on *O_i,t_*> 0. The likelihood of *λ_t_* is then

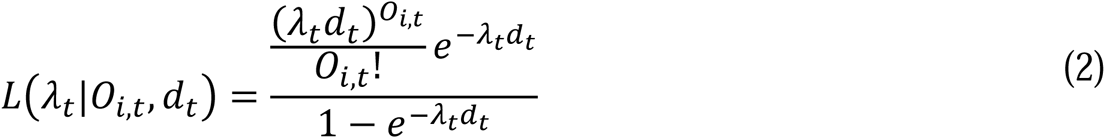

The maximum likelihood estimate for the sampling rate of a group of species in a given interval is found by maximizing the product of eq 2 over all the observed species (*N_t_*) belonging to that group;

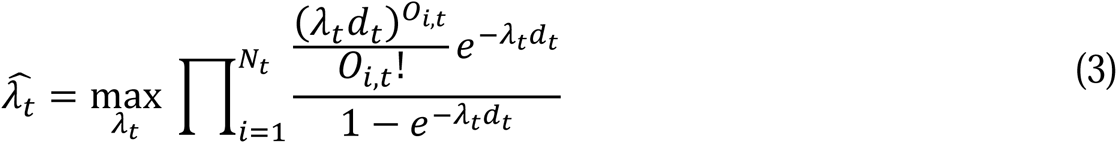

If our data consist of only single records (i.e. *O_i,t_* = 1, for all *i*), estimating *λ_t_* using maximum likelihood will yield an estimate of 0. Hence, the minimum data requirement for estimating the fossilization rate is a dataset where at least one of the species has more than one observation.

We assume that sampling rates estimated are constant for all species within a clade in the same time interval (i.e. the sampling rates estimated are time-specific but not species-specific). We can then estimate the probability of detecting a species from this group as 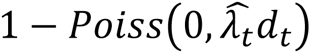, i.e. one minus the probability of not detecting a species if it was actually extant, according to the Poisson process. We further use this binomial probability in deriving the most likely true richness. The binomial probability of a species sampled during an interval *d_t_* is

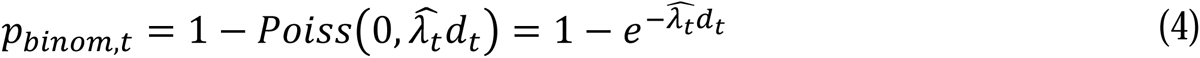

where 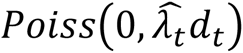 is the probability of zero sampling events in one lineage with a rate 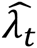 in a bin of duration *d_t_*. Note that sampling rate (*λ_t_*) and sampling probability (*p_binom_*,*_t_*) are different; while sampling rate has units observations per lineage per Ma, the sampling probability is a stage-specific probability of detecting a species that was extant during that interval. The last step in estimating the true richness in a given time interval is to find the true species richness (*N_true_*) that maximizes the binomial likelihood

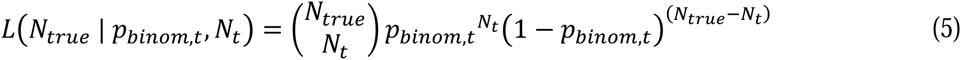

where *p_binom,t_* is the binomial probability calculated from the estimated sampling rate (eq 4) using maximum likelihood estimates (eq 3) and *N_t_* is the observed number of species in the time interval. Thus the value *N_true_* that maximizes eq 5 is the maximum likelihood estimate of the true richness where *N_t_* species were observed.

To quantify the uncertainty surrounding the estimate of the sampling rate and the true species richness we utilize the relationship between the χ^2^ distribution and log likelihood profiles (see e.g. 29). For the confidence bounds on the maximum likelihood estimate 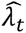 we find the range of values for *λ* that satisfy the inequality

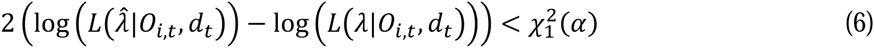

where 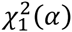 is the upper quantile function of the χ^2^ distribution with one degree of freedom. Similarly the upper and lower confidence bounds for the estimated true richness *N_true_* is found using the lower and upper confidence bounds on the sampling probability (*p_binom,t_*).

TRiPS thus yields maximum likelihood estimates and confidence intervals of true species richness for a given time interval by estimating a sampling rate (observations per species per Ma). This sampling rate can be transformed into a time interval specific sampling probability (probability of fossil detection per species) and thereby appropriately take the duration of the time interval into account. In other words, we do not need to conform the data to equal durations as commonly done (22,34). The sampling rates estimated from TRiPS are thus directly comparable across geological intervals of unequal durations. Note that while we have described TRiPS using species observations it can also be directly applied to genera or groups of taxa defined in other ways. In fact, any grouping of taxa thought to exhibit similar sampling rates might be combined, whether or not they actually are taxonomic clades.

### Simulations using a birth-death-fossilization process

To evaluate our method’s applicability and power we performed a large number of continuous time birth-death (BD) simulations, coupled with a fossilization scheme, which we interpret as sampling. In a classic BD process a lineage either gives rise to a new species or goes extinct at a certain rate; our fossilization scheme adds a third potential event: that of a lineage leaving a fossil. We are thus simulating a ‘fossil record’ given a set of parameters controlling the dynamics and sampling of the simulated clade, and then using TRiPS to estimate the true number of species in these simulations

In case of no changes in species richness within a given time interval and identical sampling rates for all lineages, TRiPS will consistently recover true richness. Using simulations, we explicitly investigate the robustness of our approach to violations of TRiPS’s two main assumptions, 1) equal sampling rates (per lineage per Ma) for all species in the clade in question and 2) and negligible species turnover within a time interval (i.e. all lineages span the entire interval in which they are observed). We then use the results of our simulations to aid interpretation of our estimates based on dinosaur records (see below).

Our birth-death-fossilize model has six parameters; speciation and extinction rates (in per species per time unit), the number of species at the start of the simulation, the duration of time units (in continuous time), mean sampling rate (fossils per species per time unit), and a parameter that scales the variability of sampling rates among individual species.

For our simulations, speciation and extinction rates were drawn from log-uniform distributions spanning 0.001 to 0.15 per species per time unit. For comparison, the maximum mean speciation rate for birds in the Cenozoic has been estimated to be approximately 0.15 (35). Note that having non-zero speciation and extinction rates allow for changes in species richness, which explicitly violates the assumption of all lineages spanning the whole interval in which they are found. The initial number of species in the simulations ranged from 10 to 250, drawn from a uniform distribution. Durations of time units simulated ranged from 2 to 20, drawn from a uniform distribution mimicking the durations of the geological stages in the Mesozoic where the Hettangian (2 Ma) and Norian (19.5 Ma) are the shortest and longest stages, respectively. Mean sampling rates ranged from 0.001 to 1.5 fossils per species per time unit as drawn from a uniform distribution. Variation in sampling rate among species ranged from 0 to 0.3, also drawn from a uniform distribution; setting this parameter to 0.3, for example, gives each lineage a sampling rate ranging from 0.7 to 1.3 times the clade mean (this is equivalent to twice the coefficient of variation). Each species thus has its own unique sampling rate drawn from a uniform distribution with means and variance differing across simulations, explicitly violating the TRiPS assumption that lineages have the same sampling rate. We ran 100 000 simulations.

We evaluated TRiPS’s ability to infer sampling rates and true species richness by 1) tabulating the number of simulations in which the true species richness was inside the predicted confidence interval (which we call success rate), 2) estimating the bias in the maximum likelihood prediction of species richness, the mean scaled error 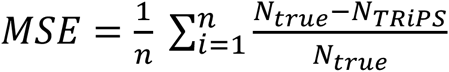, and 3) Pearson’s product-moment correlation (*ρ*) between true and estimated richness in the simulations. We compiled the same statistics for estimated sampling rates and sampling probabilities. Together these three metrics inform us of the precision, bias and relative accuracy of our methodology.

### Estimating dinosaur sampling bias and species richness

For each stage in the Mesozoic we estimate the sampling rate (i.e. the bias) and the species richness for each of the four clades using TRiPS. Combined with confidence intervals, these estimates allow for sound statistical comparisons of both bias and richness between geological stages. Additionally, to give a crude estimate of the total richness across the Mesozoic, we calculate a Mesozoic sampling probability as a mean of binomial probabilities, weighted by estimated richness in each stage. Mesozoic mean sampling rates are calculated similarly. Using mean Mesozoic sampling probability and the number of unique observed species we could also estimate the total richness of the dinosaur clades throughout the Mesozoic.

### Implementation

All data analysis was performed in *R* (36). Code necessary for the analysis, combined with scripts to directly download relevant (and thus updated) data from Paleobiology Database in addition to simpler R scripts showing how other datasets can be analysed using TRiPS is available on the authors’ website and Dryad. Output from our simulations and functions and scripts to rerun simulations are also deposited,.[AUTHOR COMMENT: Dryad will not accept data before acceptance of manuscript].

## Results

### Dinosaur sampling estimates among clades and through time

Ornithischian species have the highest Mesozoic mean sampling rate (0.241 sampling events per lineage per Ma (0.172 – 0.355), see table 1). Sauropodomorph and theropod species have similar mean sampling rates with 0.156 (0.084 – 0.304) and 0.153 (0.104 – 0.243) respectively, and all dinosaurs have on average been sampled 0.192 (0.146 – 0.262) times per species per Ma. Mesozoic mean species sampling probabilities are also different among clades; theropods (0.463; 0.335 – 0.628) and sauropodomorphs (0.479; 0.278 – 0.753) have lower sampling probabilities than ornithischians (0.708; 0.575 – 0.847).

**Table 1.** Mesozoic mean sampling rates (observations per lineage per Ma), sampling probability and total richness for each of the four clades, with 95% confidence intervals in parentheses. Note that the mean rates and probabilities were calculated weighted by the estimated richness (see ESM, table S2). Total Mesozoic richness was calculated using the Mesozoic mean sampling probability and the total number of species in each clade. For stage specific sampling estimates see figure 1 and ESM.

| Clade | Mesozoic sampling<br>rate | Mesozoic sampling<br>probability | Total Mesozoic<br>species richness |
| --- | --- | --- | --- |
| **Dinosauria** | 0.192 (0.146 – 0.262) | 0.580 (0.474 – 0.706) | 1936 (1543 – 2468) |
| **Ornithischia** | 0.241 (0.172 – 0.355) | 0.708 (0.575 – 0.847) | 508 (409 – 668) |
| **Sauropodomorpha** | 0.156 (0.084 – 0.304) | 0.479 (0.278 – 0.753) | 513 (307 – 983) |
| **Theropoda** | 0.153 (0.103 – 0.245) | 0.463 (0.335 – 0.628) | 1115 (780 – 1653) |

TRiPS based stage-specific sampling rates and probabilities for dinosaurs do not monotonically increase through the Mesozoic, but exhibit a combination of high and low sampling regimes (figure 1). This observation runs counter to the commonly held belief that younger geological strata exhibit a higher level of fossil sampling (e.g. 21,23). Sampling probabilities are particularly high during the Hettangian (201.3 – 199.3 Ma) and Sinemurian (199.3-190.8 Ma), the Tithonian (152.1-145 Ma), the Albian (113-100.5 Ma) and the Maastrichtian (72.1-66 Ma) but these high sampling intervals are interspersed with lower ones. Note that sampling rates (*λ*, top panel figure 1) and sampling probabilities (*p_binom,t_*, lower panel figure 1) while showing some commonalities, are not the same. For instance, the Norian (228-202.5 Ma) has relatively few sampling events per lineage per million years, but the probability of a species being sampled, given that it was extant, is quite high (>0.8 for all groups, see figure 1). In general, the relative changes in sampling dynamics are similar for our genus level analyses although sampling rates and probabilities are naturally higher for genera (ESM figures S5 and S6).

**Figure 1.**
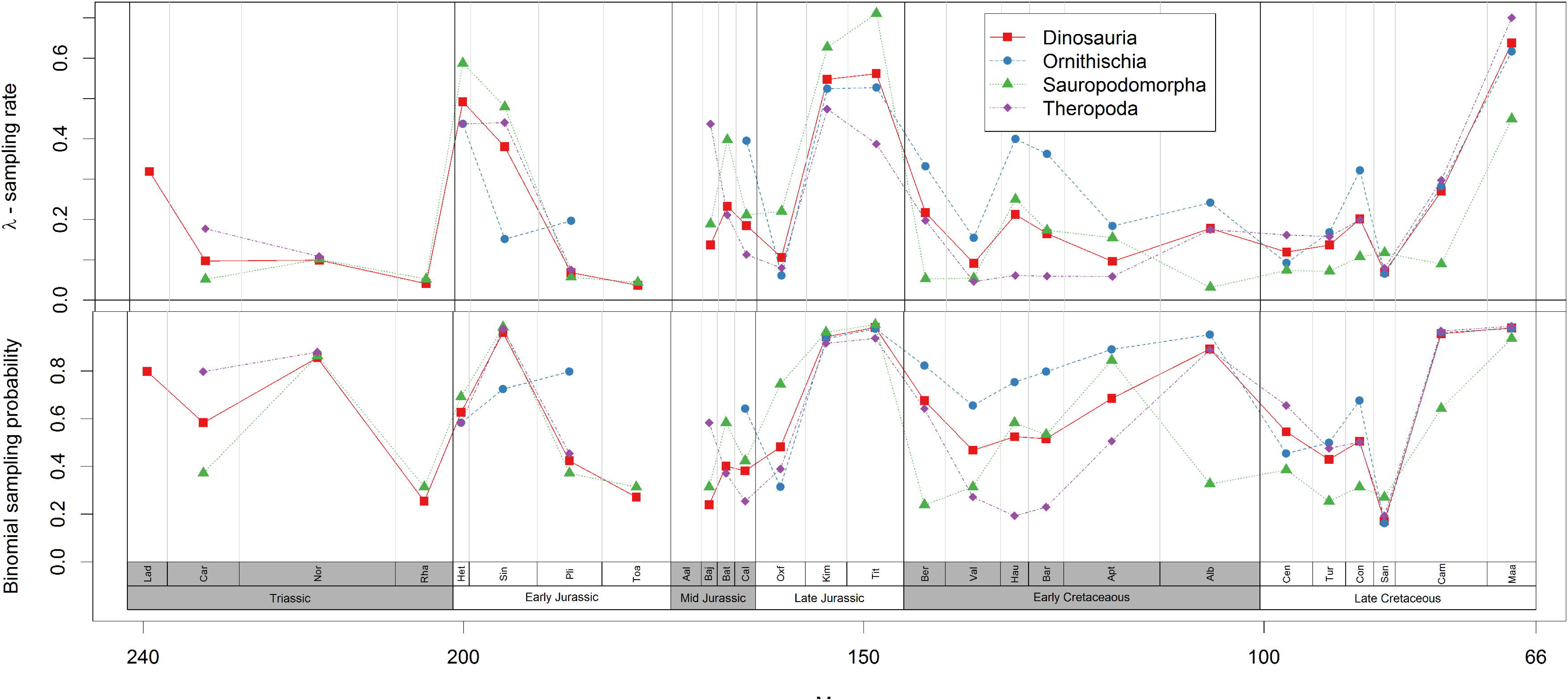
<u>Species sampling estimates from TRiPS for all dinosaurs, ornithischian, sauropodomorph and theropod</u>. Top: Estimated sampling rates (*λ_t_*) in sampling events per species per million years. Bottom: Estimated binomial sampling probabilities (*p_binom,t_*) of species within the plotted time interval. In both panels, estimates are in red (squares and full line) for all dinosaurs, blue for Ornithischia (circle and dashed line), green for Sauropodomorpha (triangles with dotted line) and purple for Theropoda (diamonds with dash-dotted line). Confidence intervals on all rates and probabilities and estimates from 100 replicated observation count matrices (see main text) are reported in the Electronic Supplementary Information. Abbreviations used: Lad – Ladinian, Car – Carnian, Nor – Norian, Rha – Rhaetian, Het – Hettangian, Sin – Sinemurian, Pli – Pliensbachian, Toa – Toarcian, Aal – Aalenian, Baj – Bajacian, Bat – Bathonian, Cal – Callovian, Oxf – Oxfordian, Kim – Kimmeridgian, Tit – Tithonian, Ber – Berriasian, Val – Valanginian, Hau – Hauterivian, Bar – Barremian, Apt – Aptian, Alb – Albian, Cen – Cenomanian, Tur – Turonian, Con – Coniacian, San – Santonian, Cam – Campanian, Maa – Maastrichtian.

The three clades have notably different sampling estimates from stage to stage, with binomial sampling probabilities spanning from about 0.1 to almost 1. Theropods show higher sampling rates relative to ornithischians and sauropodomorphs in the Triassic (Carnian, 237 – 227 Ma, and Norian) but comparably lower rates in parts of the Early Cretaceous (Valanginian, 139.8 – 132.9, through Aptian, 125 – 112 Ma). This runs counter to earlier conclusions that richness trajectories of Theropoda and Ornithischia seems to largely be driven by sampling bias, whereas Sauropodomorpha are less affected by bias in the fossil record (25,26).

The different sampling rates across stages give a very different picture of bias than what the residual approach does. To reiterate, the residual approach assumes that a chosen time series fully captures the sampling bias, and uses a model of fixed diversity to predict how richness would look *if only* biased sampling drove the detected signal (8,9). The number of fossil collections from different intervals that contains at least one dinosaur (DBCs) are often used as a sampling proxy for dinosaurs (23,26,37). We compare our proxy-free sampling estimates to DBCs to check how much sampling bias DBCs capture. We performed Pearson correlation tests of the binomial sampling probabilities estimated and the linearly detrended log10 number of collections for all downloaded dinosaur observations (see ESM table S2). These estimates indicate that there is a common signal in our estimated sampling probabilities and DBCs with correlations of 0.50 (p=0.0090), 0.64 (p=0.003), 0.71 (p= 9e-5) and 0.60 (p=0.002), for Dinosauria, Ornithischia, Sauropodomorpha and Theropoda, respectively. However, there is substantial remaining sampling bias not captured by DBCs (since R^2^ < 0.51 for all). Using DBCs to remove bias in richness estimation of dinosaurs (e.g. 38) will add a substantial amount of error in correcting diversity curves.

### Dinosaur richness in the Mesozoic

The whole Mesozoic is estimated to have seen 1936 (1543 – 2468, table 1) dinosaur species. A Mesozoic mean sampling probability at the genus level (again, weighted by estimated genus richness from each stage) yields estimates of total dinosaur genera richness at 1536 (1255 – 1929). Total species richness for the subclades is estimated to be 508 (409 – 668) for Ornithischia, 513 (307 – 983) for Sauropodomorpha and 1115 (780 – 1653) for Theropoda (table 1).

The species richness estimates from TRiPS shares dynamics with those painted by both the raw counts of species and range-through species richness using the same dataset (figure 2). However, only in about half the stages are the range-through estimates within the confidence interval of TRiPS estimates. Genus richness dynamics are similar to species dynamics (ESM figure S6) and indicate that for at least this dataset using these analyses, genus level estimates can be a proxy for species estimates, corroborating Jablonski and Finarelli’s findings (39). While genus richness estimates are lower, they are similar to species estimates, unsurprisingly, given there are few dinosaur species per genera (approximately 1.15 identified species per genera in our data).

**Figure 2.**
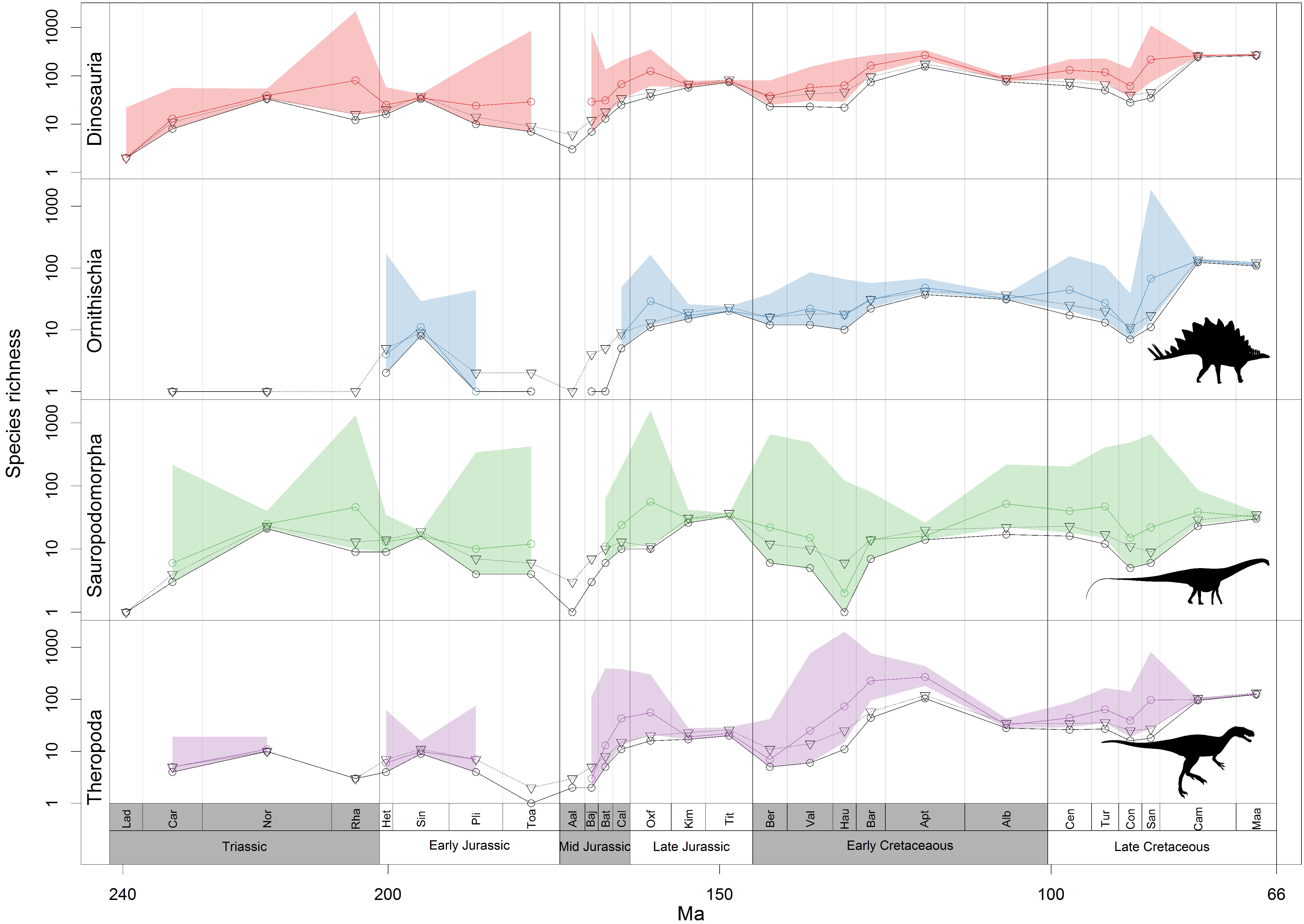
<u>Species richness estimates from TRiPS</u>. Black circles connected by full line indicate observed species counts, triangles connected by dotted line indicate range-through species counts while coloured line and shading indicate maximum likelihood estimate and 95% confidence intervals for true species richness estimated using TRiPS. Corresponding sampling estimates can be seen in figure 1. All estimates with confidence intervals are also in the Electronic Supplementary Information.

Estimated species richness shows an increase in the late Triassic, with a peak in the Rhaetian, though with a considerably wide confidence interval (figure 2, ESM table S2). This indicates that the initial diversification of dinosaur continued for up to 40 Ma, in contrast to the raw counts which peak in the Norian. Comparing stages within the Early Jurassic shows that the elevated raw richness in the Sinemurian for all clades is not corroborated by our estimates; TRiPS estimate richness for this stage well within the confidence intervals for richness in the Hettangian, Pliensbachian (190.8 – 182.7 Ma) and the Toarcian (182.7 – 174.1 Ma). In the Mid and Late Jurassic, also contrary to the raw species counts indicating a richness peak in the Tithonian, TRiPS estimates that all clades exhibit elevated richness in the Oxfordian due to the relatively low sampling rate in this interval for all clades. Finally, the trough in raw richness in the Coniacian (89.8 – 86.3 Ma) - Santonian with a peak in the two final stages of the Cretaceous for all clades seems to be a sampling artefact; while there is still a signal of reduced richness in the Coniacian for all dinosaurs, this is most likely driven by the ornithischians; the Coniacian-Santonian trough is not particularly strong for sauropodomorphs and theropods when inspecting the confidence intervals for these periods.

### Simulation results

Most simulated species do not span the whole interval in which they were sampled, unless speciation and extinction rates are both zero. Using TRiPS to estimate the true richness is in such cases (the majority of our simulations) is thus a test of the degree to which our approach is robust to deviations from the assumption that an observed lineage spans the whole interval in question. Across the whole parameter space simulated (see above) TRiPS yielded confidence intervals including the true richness in 36% of simulations and true sampling rates were inside the confidence interval in 39% of simulations (see ESM for details). Though these are low absolute success rates, the correlations between true and estimated richness is still high (*ρ* = 0.99) and the mean scaled error is also low (MSE<0.07). This indicates that the maximum likelihood estimate of richness correlates very strongly with the true richness (indicating robustness in terms of *relative* richness) and that the richness estimate is, on average, only 7% smaller than the true richness. Unsurprisingly, the success rate and MSE of TRiPS are also highly dependent on all parameter settings, except the degree of variation in sampling rate between lineages in a single simulation (see ESM). In cases where individual lineages differ in sampling rates, TRiPS effectively estimates the group mean, and there is no first order effect on bias, precision or correlation of estimates when increasing the variability of sampling rates between lineages. The duration of the interval directly scales the potential for changes in species richness; the longer the duration, the more time for speciation and extinction within the time interval, leading to larger deviations from our assumption of lineages spanning the entire interval, and a higher failure rate of capturing the true richness within the confidence interval predicted by TRiPS. This underscores that TRiPS will perform better when applied to empirical data with narrower time intervals. Secondly, the sampling rate itself affect the success rate of TRiPS; TRiPS works best where sampling rates are relatively low, and even so when speciation and extinction rates are > 0 (figure 3). We note, however, that the majority of our empirical dinosaur estimates are in regions of parameter space in which TRiPS has a relatively high success rate (figure 3).

**Figure 3:**
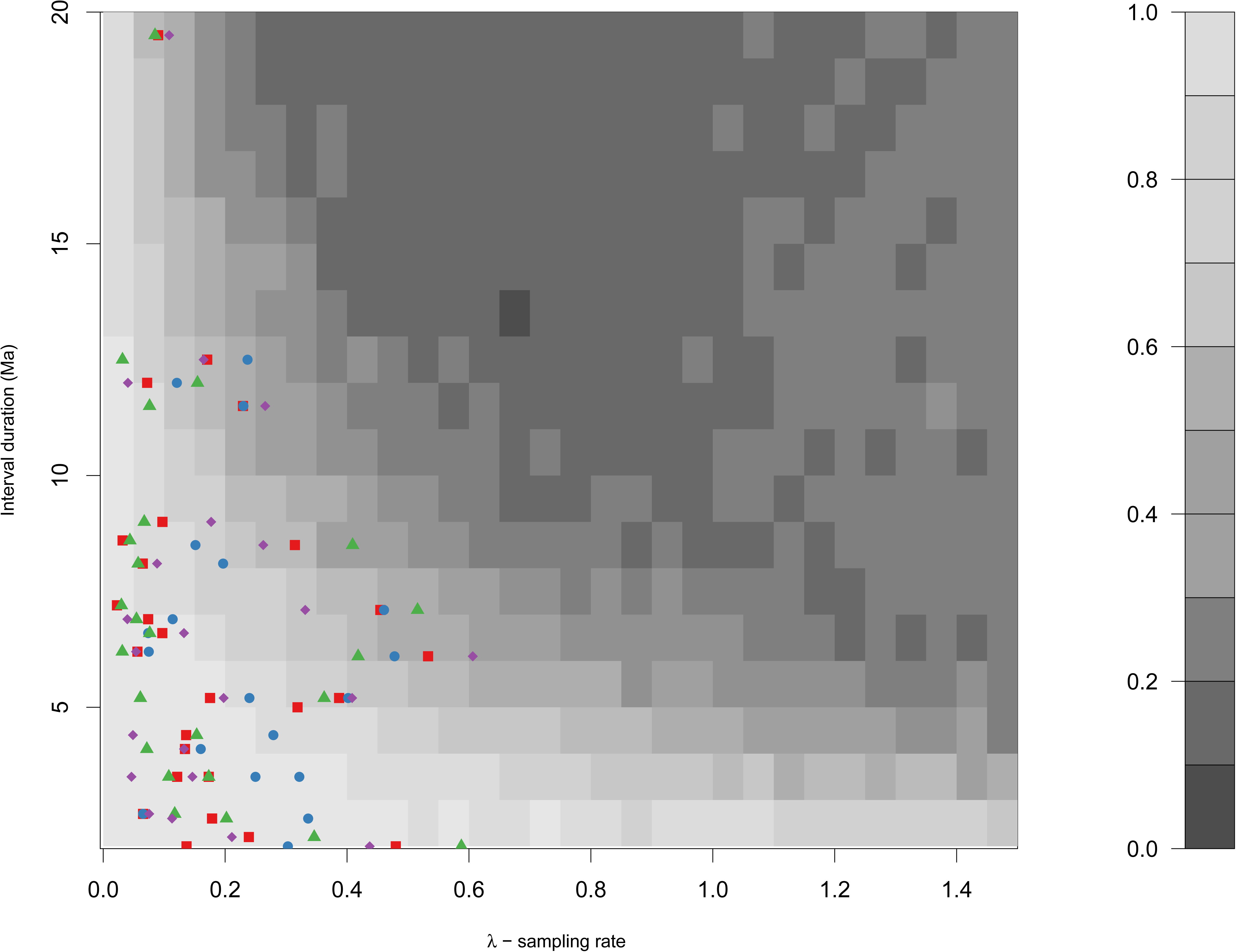
<u>Simulation and estimation results.</u>. The grayscale indicates TRiPS’s success rate (the fraction of simulations that had confidence intervals that span the true value) in the simulations, lighter areas indicate higher success rate. For each combination of duration and sampling rate in the figure above, the full range of all other parameters are represented (i.e. the success rate is the mean value over sampled speciation and extinction rates, as well degree of individual lineage variability in sampling rate and number of initial lineages, see main text). Estimated sampling rates for Dinosauria (red squares), Ornithischia (blue circles), Sauropodomorpha (green triangles), Theropoda (purple diamonds) are plotted against their corresponding stage durations.

## Discussion

Fossil observation data are readily available in public databases, such as the PaleoDB. Yet, estimating taxon richness using such databases is not trivial as fossilization, outcrop exposure and modern day sampling and data compilation are heterogeneous processes. Unlike approaches such as subsampling (1,34,40) and bias-corrected residual analysis (8,9) our approach, TRiPS, estimates true rather than relative richness by utilizing information on sampling which is inherent in PaleoDB. In addition, unlike the residual approach, we do not make presuppositions that an external time series can be used to correct for sampling. This is important because such external time series (e.g. amount of outcrop, sea level) may constitute a factor driving both richness and sampling as postulated by the Common Cause Hypothesis (10,11) or be an effect of a third factor. The residual approach can also suffer from redundancy in the data used (41,42), which would lead to erroneously corrected richness estimates.

In TRiPS, we tackle bias in the fossil record directly by estimating rates of sampling. This also allows us to disentangle sampling and richness dynamics such that tests of links between potential drivers can be done on sampling and richness independently (see also 43). An advantage of TRiPS is that our treatment of sampling allows sampling probabilities to be directly comparable between intervals of unequal duration (but see below). One assumption we do explicitly make which cannot be true most of the time, is that a species detected in a given time interval is extant during that whole time interval. This is because most species are unlikely to become extinct exactly at the late boundary of a time interval or originate exactly at the early boundary of a time interval. While other methods for estimating richness also assume that turnover is clumped at interval boundaries (see e.g. 40, p. 74), we explicitly examined the robustness of our estimates to the violation of this crucial assumption.

Comparing our empirical sampling rate estimates with simulations that violated key assumptions of TRiPS (figure 3), we find most of our empirical estimates fall within parameter ranges in which we are able to retrieve true richness estimates reliably. This is with the caveat that the simulated speciation and extinction rates are realistic for dinosaurs.

However, we note other caveats to the estimates from TRiPS. First, although the ability to estimate true richness is relatively robust to deviations from our assumptions under our simulations, TRiPS does give biased estimates of both sampling and richness when there are within-bin dynamics. When applying TRiPS to cases with non-zero speciation and extinction rates and increasingly wide time intervals, our ‘data’ will consist more and more of observations of lineages that did not span the entire interval. This generates a bias in our estimated sampling rates such that these estimates are lower than true sampling rates (see ESM figure S2), and more so for longer durations and higher levels of speciation and extinction rates. On the other hand, transforming this rate into a binomial probability (eq 4), assuming all lineages lasts longer than they actually do, leads to an overestimation of the binomial sampling probability when compared to the fraction of species actually observed in our simulations (see ESM figure S3). These two opposing biases in our methodology in sum lead to a slight negative bias in the reconstructed richness (on average 7% below true richness in our simulations). So, sampling rates and richness are slightly underestimated, but sampling probabilities are overestimated.

The estimated richness is thus best treated as minimum richness estimates, particularly for intervals in which there is reason to believe that within interval changes in true richness have been substantial, such as in long geological stages. Second, with longer intervals (which gives more time for in-bin dynamics) and higher sampling rates TRiPS fails more often in terms of capturing true richness inside confidence intervals (figure 3) due to high precision combined with bias. Note, however, that the maximum likelihood estimate of true richness itself is still not too far off target if speciation and extinction rates are not too high (see ESM figure S1). On the other hand, one benefit of our explicit approach is that it is straightforward to simulate a birth-death-fossilization process to check if the empirical estimates of sampling rates and richness can be considered robust to violation of the assumption of negligible turnover within an interval (see below). It is also worth highlighting that other approaches for reconstructing past richness also fall victim to deviations from constant richness (see e.g. 40), even though such violations have not been explicitly examined in published simulations, as far as we know.

The obvious solution to these caveats is to apply TRiPS only for temporally well-resolved data, and to strive for better and more accurate dating of fossils. Other potential avenues for improvement lie in development of the methodology itself. For instance, extending TRiPS to estimate sampling rates in a hierarchical manner, where instead of estimating a single sampling rate, one could estimate a distribution of sampling rates within a time interval, or link sampling rates for subclades in a hierarchical fashion, possibly including more phylogenetic information. Other directions could be to include estimates of lineage durations from independent sources instead of assuming that all lineages span the interval in which they are observed.

In summary our simulations shows that TRiPS is prone to errors if its assumptions are substantially violated. Even in applications where we have reason to believe there are high levels of species turnover, estimates of richness will still be a reasonable approximation of the true richness, although the associated confidence intervals should be treated with caution; they are in general too narrow for such cases. In applying TRiPS on shorter intervals, i.e. when there is more reason to believe that lineages sampled actually do span most of the interval, TRiPS confidence intervals will in most cases encompass the true richness. With the pros and cons of TRiPS in mind, we can now compare our dinosaur estimates to previous studies.

Dodson (19) estimated the total number of dinosaur genera to be 900-1200 for the whole Mesozoic, with about 100 genera at any one geological stage. Our estimated genus richness of 1536 (1255 – 1929) are more in line with Wang and Dodson’s (18) estimates, which inferred that the entire Mesozoic saw 1844 dinosaur genera. For the final stages of the Cretaceous Wang and Dodson (18) estimated 200-300 genera of dinosaurs roaming our planet, corroborated by our estimates of 221 and 227 genera for Campanian and Maastrichtian respectively (see ESM figure S5, ESM table S2).

It is currently accepted that dinosaurs did not rapidly diversify when they appeared around the start of the Late Triassic. Rather, sauropodomorphs diversified during the final part of the Triassic, while ornithischians and theropods increased in richness in the early Jurassic (21,38). While this pattern is in part corroborated by our analysis for sauropodomorphs, a species richness of 80 for the whole dinosaur clade is estimated for the last stage in the Triassic, which is relatively high compared with the rest of the Mesozoic. Sampling rates for ornithischians cannot be estimated with confidence for any interval in the Triassic. Sauropodomorphs exhibit rather high levels of both observed and estimated species richness already in the Norian (228 – 208.5 Ma), and our estimate of sauropodomorph species richness during the Rhaetian (208.5 – 201.3 Ma) is so high that it was not even surpassed by the diversity in the final stages of the Late Cretaceous, and only barely so for the diversity peaks in the Late Jurassic (44) and Early Cretaceous. In other words, our results indicate that both sauropodomorph and dinosaur richness peaks in the Rhaetian, and not in the Norian as earlier studies (26), due to the estimated low sampling this final stage of the Triassic. This is in contrast to previous residual based analysis which indicate the Rhaetian shows a marked decrease in diversity (38, but also see 9).

It is largely accepted that the Oxfordian (163.5-157.3 Ma) exhibits remarkably low diversity; perhaps it could even be considered the most depauperate stage throughout the Age of the Dinosaurs (21,26,38,44), and particularly so for sauropodomorphs (26). Our approach, in contrast, estimates the sampling probability in this particular stage to be the culprit of this trough and especially so for sauropodomorphs (see figure 1). Instead of inferring low species richness in this stage, our estimates indicate the sauropodomorph richness has doubled from the previous stage (Callovian), and a great richness increase is also evident in ornithischians and the dinosaurs as a whole. More intense sampling efforts, both in the field and in museum collections, and detailed analysis on the observations from the Oxfordian are clearly needed.

Some have argued that the Jurassic-Cretaceous (J/K, Tithonian - Berriasian) boundary (~145 Ma) demonstrated a clear diversity loss particularly pronounced for sauropodomorphs (25,26,38,45), while others have claimed otherwise (46,47). Though the raw species counts partially corroborate a decline in richness across the J/K boundary, the sampling rates for the early part of the Cretaceous are estimated to be much lower than late Jurassic (figure 1), and the difference in sampling rates are particularly pronounced for the sauropodomorph clade. While the richness for all dinosaurs is estimated to decrease (figure 2 top panel), this signal is equivocal for any of the subclades whose confidence intervals for richness in the Berriasian are wide (and span the richness in the Tithonian) due to low and uncertain sampling probabilities. Genus level analyses (see ESM Table S2) estimate that the number of sauropodomorph genera in Berriasian is in fact the same before and after the J/K boundary. It is also worth emphasizing that the confidence intervals for the estimated species diversity are much wider in Berriasian compared to Tithonian, implying that an increase in the true richness across this boundary cannot be excluded. Lower sampling rates during the Berriasian across all dinosaur clades and regardless of whether species or genus level data are used, suggest that this “clear diversity loss”, might be an artefact of sampling bias and that the ‘major extinction’ of dinosaurs across the Jurassic-Cretaceous boundary (25,26,44,48,49) might be less severe than previously thought (see also 45,46).

There current view is that not only is there no longer-term (i.e. epoch-level) decline in richness prior to the end-Cretaceous extinction, there is in fact a sharp increase in richness for the final two stages in the Cretaceous (50). The latter is not corroborated by our estimates; while the two final stages of the Cretaceous are high in raw species counts for all clades, estimated richness in Campanian and Maastrichtian are well within confidence intervals for earlier stages in the Cretaceous for sauropodomorphs, and also partially for ornithischians and theropods. Rather than the two final stages exhibiting elevated richness levels, TRiPS estimates that similar level of richness was reached in the previous stage (Santonian), at least for Theropoda and Ornithischia. Ornithischia do seem to have a real trough in the Coniacian (89.8 – 86.3 Ma), but sauropodomorphs and theropods seem instead to have steady, but slow, decreases and increases in richness, respectively, across the Late Cretaceous.

Late Cretaceous increase in dinosaur diversity has also been framed as a debate on whether or not dinosaurs showed a decline in species richness prior to the Cretaceous-Paleogene extinction event (9,18,21,24,25,50). Brusatte et al. (50) argued that, while there was no global long-term decline prior to the end-Cretaceous extinction, there is evidence for ceratopsids and hadrosaurids (members of Ornithischia) exhibiting declines in both species richness and morphological disparity in the final 15 Ma of the Cretaceous (50). On the other hand, Lloyd (9) claimed that sauropodomorphs and ornithischians both show long-term declines throughout most of the Cretaceous (using the residual approach) whereas Barrett et al. (25) highlighted a negative trend in taxic diversity for theropods and ornithischians in the last two stages of the Cretaceous, but suggested a ‘radiation’ of sauropodomorphs in the Late Cretaceous (see also 44). TRiPS estimates the final stage of the Cretaceous to be about 13 % less speciose for ornithischians (figure 2, ESM table S2) and a reduction in species richness by 17% is estimated to have occurred for sauropodomorphs, though with less confidence, while we estimate a significantly higher richness in the Maastrichtian than the Campanian for theropods (see ESM table S2).

### Conclusions and future directions

To paint an accurate picture of past species richness and to identify of periods of high diversification and major extinction events, the bias inherent the fossil record that may mislead and confound our inferences needs to be taken into account. Here we have detailed TRiPS, a new approach for estimating both temporally varying sampling and species richness. The application of TRiPS to a global dataset of dinosaur records indicates that several of the commonly held ideas about the species richness trajectory of dinosaurs might be effects of either sampling bias or the use of methods that might have introduced new biases to richness estimates through their assumptions.

As a tool that estimates both sampling rates *and* true richness directly, TRiPS is pregnant with possibilities and have applicability to a range of other palaeontological questions. Richness and sampling estimates from TRiPS allow us to test the Common Cause Hypothesis in a straightforward manner if potential common drivers can be measured in the geological record. Estimates of sampling can be used in predicting true ranges of a given species, if we can make the assumption that species have the same temporally varying sampling rates. The two forms of sampling estimates may help palaeontologists focus their sampling and taxonomic efforts in time intervals (or geographic regions) which require most effort given the specific questions we wish to answer. Additionally, sampling rates are needed for better calibration of phylogenetic trees using fossil observations, and for inference of rates of speciation and extinction using phylogenies on extinct taxa. While the application of TRiPS we demonstrated here attempts to estimate global richness of dinosaurs and its major subclades, TRiPS can be applied to any collection of lineages that are assumed or shown to have similar sampling rates, and could also be used to estimate taxonomic richness on local to continental scales.

### Data accessibility statement

All data were downloaded from Paleobiology Database (https://paleobiodb.org/#/). A copy of the download is available on Dryad [ref to Dryad here].

### Authors’ contributions

JS conceived the study, performed the analysis and wrote the manuscript with substantial contributions to writing and interpretation by LHL. Both authors gave final approval for publication.

## Acknowledgements

Kjetil Lysne Voje is acknowledged for fruitful discussions surrounding the initial idea, and Fakutsi is recognized for inspiration. We are also thankful for comments and criticisms made by Philip Mannion and Manabu Sakamoto acting as reviewers. Comments on the manuscript and ideas inside it by Roger Benson, Matthew Clapham, Gene Hunt, Trond Reitan and Steve Wang are also acknowledged. We also thank Benson for sharing his lists of ichno- and ootaxa. Credits to Andrew A. Farke, Scott Hartman and Conty, via PhyloPic, for the silhouettes in figure 2. This work would not be possible without vertebrate palaeontologists who contributed primary data and the authorizers and enterers of the PaleoDB who compiled these primary data. Major contributors to the downloaded PaleoDB data are Matthew Carrano (59,8%), John Alroy (13,9%), Mark Uhen (8,9%), Anna Behrensmeyer (4,6%) and Philip Mannion (3,1%). This is PaleoDB Publication XXX.

## Electronic supplementary material (ESM)

Electronic supplementary material contains details of the simulations, results regarding these simulations,, results from estimating sampling from all 100 replicated datasets detailed in the main text. An Excel file (ESM_Table_S2.xlsx) contains all estimated sampling rates, sampling probabilities and true richness for each clade in each stage at both species and genus level. Data and R code to replicate analysis and simulations is available on Dryad [ref to dryad here]

